# Tyrosine transfer RNA levels and modifications during blood-feeding and vitellogenesis in the mosquito, *Aedes aegypti*

**DOI:** 10.1101/2023.11.29.569187

**Authors:** Melissa Kelley, Christopher J. Holmes, Cassandra Herbert, Asif Rayhan, Judd Joves, Melissa Uhran, Ronja Frigard, Khwahish Singh, Patrick A. Limbach, Balasubrahmanyam Addepalli, Joshua B. Benoit

## Abstract

Mosquitoes such as *Aedes aegypti* must consume a blood meal for the nutrients necessary for egg production. Several transcriptome and proteome changes occur post blood meal that likely corresponds with codon usage alterations. Transfer RNA (tRNA) is the adapter molecule that reads messenger RNA (mRNA) codons to add the appropriate amino acid during protein synthesis. Chemical modifications to tRNA enhance codons’ decoding, improving the accuracy and efficiency of protein synthesis. Here, we examined tRNA modifications and transcripts associated with the blood meal and subsequent periods of vitellogenesis in *A. aegypti*. More specifically, we assessed tRNA transcript abundance and modification levels in the fat body at critical times post blood-feeding. Based on a combination of alternative codon usage and identification of particular modifications, we identified that increased transcription of tyrosine tRNAs is likely critical during the synthesis of egg yolk proteins in the fat body following a blood meal. Altogether, changes in both the abundance and modification of tRNA are essential factors in the process of vitellogenin production after blood-feeding in mosquitoes.

## Introduction

The yellow fever mosquito *Aedes aegypti* is a vector for many viruses that impact approximately 700 million people annually (Gaythorpe et al., 2021; Piovezan-Borges et al., 2022). Due to the global implications associated with this vector (Messina et al., 2019), there is a constant need for novel targets to manage mosquito populations. In anautogenous mosquitoes, females must consume a blood meal to provide the resources for egg production and associated reproductive processes, such as vitellogenesis (Attardo et al., 2005; Hansen et al., 2014). Given the availability of a blood meal is often unpredictable, a pre-vitellogenic arrest period regulated by hormones is required for preparation to allow reproduction to commence immediately after a blood meal (Attardo et al., 2003; Valzania et al., 2019; Zhu et al., 2000). Following blood ingestion, vitellogenin is synthesized in the fat body and transported to the ovaries for oocyte generation (Attardo et al., 2003; Kokoza et al., 2001). Blood digestion also promotes protein synthesis through trypsin and other proteases to break down the protein in the blood meal (Brandon et al., 2008; Nuss and Gulia-Nuss, 2023). Along with serving as a food source, consuming a blood meal induces stress responses such as heat shock and oxidative stress response proteins (Benoit et al., 2011; Benoit and Denlinger, 2017; Bottino-Rojas et al., 2019). Altogether, multiple molecular processes are involved during blood-feeding and processes underlying reproduction in mosquitoes that require substantial changes in protein synthesis.

During protein synthesis, transfer RNA (tRNA) is required to decode the messenger RNA (mRNA) codons and incorporate the appropriate amino acid into the peptide chain. The levels of specific tRNAs and their biochemical properties directly impact the decoding rate of mRNA. Chemical modifications to tRNA are abundant and necessary to tRNA structure and function (Agris et al., 2007). Modifications located on the anticodon affect the decoding of codons as well as the efficiency and speed of protein synthesis (Agris et al., 2007). Anticodon modifications are often reflected in codon usage bias in abundant transcripts and stress-specific transcripts (Chan et al., 2018; Endres et al., 2015). Similarly, transcript levels of mature tRNA correlate with codon usage and are altered in response to various physiological changes (Torrent et al., 2018). Modifications and tRNA transcript levels are known to be affected by stress (Chan et al., 2012; Torrent et al., 2018), tissue type (Huang et al., 2021), and physiological states (i.e., sex, growth phase) (Eng et al., 2018; Kelley et al., 2022b; Rak et al., 2021). Therefore, the molecular alterations post blood-feeding likely require tuning of tRNA to account for the stress and shift to a higher expression level of reproductive transcripts.

Recently, tRNA-derived fragments have been shown to be involved in blood meal response and are sex-associated in *A. aegypti* (Eng et al., 2018). Similarly, we have demonstrated that tRNA modifications and transcripts of tRNA-modifying enzymes are sex-associated in mosquitoes (Kelley et al., 2022b). Across multiple mosquito species, female mosquitoes had higher levels of tRNA modification than males (Kelley et al., 2022b). Altogether, these studies demonstrate differences in tRNA regulation and modification based on sex, which implies there may be underlying roles for change in tRNA levels and modifications in mosquito reproduction. Given the induction of vitellogenin synthesis post blood-feeding, changes in tRNAs are likely critical to produce proteins required for mosquito reproduction. Furthermore, the elevation of multiple stress proteins post blood-feeding suggests that the expression levels and modifications of tRNAs could be involved in the stress induced by thermal, osmotic, and nutrient changes following a blood meal (Benoit et al., 2019, 2011; Bottino-Rojas et al., 2019; Lahondère et al., 2017; Sterkel et al., 2017). Thus far, it is unclear how blood meal consumption and the vitellogenic stages could affect tRNA levels and modifications in mosquitoes.

Here, we demonstrate blood-feeding status affects tRNA expression and modification levels in *A. aegypti*. Profiling of whole body samples revealed that modifications and tRNA-modifying enzyme expression are altered in a time-dependent response after blood-feeding. Furthermore, mature tRNA and precursor tRNA (pretRNA) transcripts are altered post blood-feeding, likely to meet changes in codon demand of reproductive transcripts. More specifically, tyrosine (Tyr) codons are enriched in vitellogenin and Tyr tRNA are elevated in the fat body at peak hours of vitellogenesis. Notably, recently transcribed precursor Tyr tRNAs are elevated with some tRNAs only being expressed post blood-feeding. This highlights a rapid tyrosine tRNA synthesis following a blood meal. Wobble modifications associated with Tyr tRNAs did not increase in the fat body, suggesting increases in overall Tyr tRNA levels, irrespective of anticodon modification status, are likely critical for vitellogenesis.

## Methods

### Mosquito rearing and blood-feeding

Mosquitoes were reared as previously described (Hagan et al., 2018; Holmes et al., 2022). Mosquitoes were held at 24-28°C with relative humidity between 70-80% and 12-hour light-dark cycles. Larvae were fed a mixture of ground fish food (Tetramin) and 1% yeast extract (Sigma-Aldrich). Adult mosquitoes have access to water and 10% sucrose solutions. Female mosquitoes were 5-7 days post emergence at the time of experiments. Non-blood-fed (NBF) females were not offered a blood meal. A human volunteer provided a blood meal to female mosquitoes for 30 minutes and mosquitoes were visually inspected to confirm blood ingestion. Human feeding studies were approved by the University of Cincinnati Institutional Review Board, IRB - 2021-0971 (Ajayi et al., 2021). After blood-feeding, 60-70 mosquitoes were collected at each time point (6H, 12H, 24H, 48H, 72H) and immediately frozen at -80 °C until RNA isolation.

### Expressional analyses of tRNA-modifying genes in relation to blood-feeding and reproduction

Microarray data to assess enzyme expression was collected from the GEOR2 project GSE22339 (Dissanayake et al., 2010). Three replicates for each time point consist of NBF, 3H, 12H, 24H, 48H, and 72H. The genes of interest, their homologous match, and microarray sample ID are listed in **Supp Table 1**. The enzymes were selected based on tRNA modification detected in mosquitoes and enzymes previously identified in *A. aegypti* (Kelley et al., 2022b). Each transcript has two microarray measurements; therefore, fold change was calculated using the average of the microarray expression values and with respect to non-bloodfed values. Significant differences were determined as outlined in the statistics section below.

**Table 1.** Modifications and transcript levels of modifying enzymes that have significant interactions based on time and treatment (blood-feeding status). Interactions were determined using a multivariate multiple regression model. Outputs for all enzyme modification combinations are listed in Supp Table 3.

| Modification | Gene | Treatment |  | Time |  |
| --- | --- | --- | --- | --- | --- |
|  |  | F-statistic | P-value | F-statistic | P-value |
| Ψ | PUS1 | 70.395 | 5.38E-07 | 6.882 | 0.0001 |
|  | PUS7 | 18.03 | 0.0003 | 6.871 | 0.0001 |
| D | DUS2 | 9.57 | 0.003 | 6.1 | 0.0002 |
|  | DUS3L | 5.46 | 0.02 | 4.85 | 0.001 |
| m3C | METTL6 | 7.34 | 0.009 | 6.24 | 0.0002 |
|  | METTL2B | 1.9 | 0.195 | 5.8 | 0.0003 |
| Cm | FTSJ1A | 0.285 | 0.756 | 2.42 | 0.044 |
|  | FTSJ1B | 6.8 | 0.01 | 4.98 | 0.001 |
|  | TRMT13 | 0.15 | 0.862 | 2.99 | 0.01 |
| m1A | TRMT10C | 7.02 | 0.01 | 2.86 | 0.021 |
|  | TRMT10A | 0.639 | 0.546 | 2.88 | 0.02 |
| m2G | TRMT11 | 1.63 | 0.239 | 3.14 | 0.01 |
| Gm | FTSJ1A | 1.83 | 0.205 | 1.32 | 0.278 |
|  | FTSJ1B | 20.37 | 0.0001 | 2.82 | 0.023 |
| m1G | TRMT10C | 36.96 | 1.31E-05 | 5.32 | 0.0006 |
|  | TRMT10A | 21.86 | 0.0001 | 4.97 | 0.001 |
| Q | QTRT1 | 0.037 | 0.963 | 1.92 | 0.102 |
|  | QTRT2 | 0.379 | 0.693 | 4.564 | 0.001 |
| t6A | OSGEPL1 | 12.223 | 0.001 | 1.331 | 0.275 |
|  | TP53RK | 15.54 | 0.0006 | 1.59 | 0.179 |

### Liquid chromatography tandem mass spectrometry (LC-MS/MS) for relative quantification of whole-body tRNA modifications

Isolation and purification of tRNA were carried out as previously described (Kelley et al., 2022b). Briefly, total RNA was isolated by following standard Trizol protocols. Approximately 40 whole-body mosquitoes were used per biological replicate. tRNA was separated from total RNA using a series of salt buffers and a Nucleobond anion exchange (AX100) column (Machery-Nagel, Germany) (Kelley et al., 2022b). A 1% agarose gel was performed to confirm the purity of tRNA before downstream analysis (**Supp Figure 1**).

**Figure 1:**
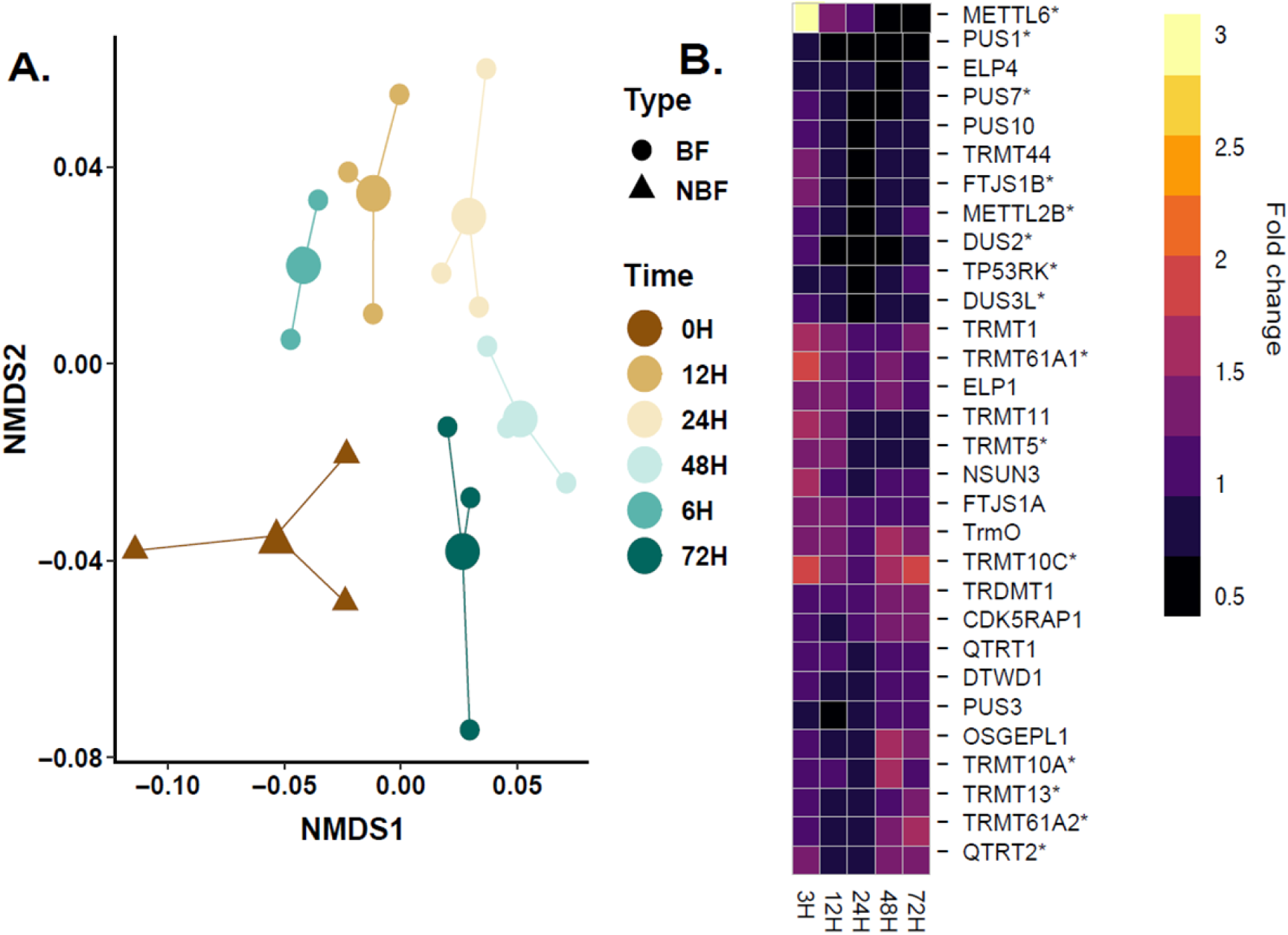
Transcripts of tRNA-modifying enzymes are affected post blood-feeding in mosquitoes. **A.** NMDS plot of tRNA-modifying enzyme expression patterns differ between blood-fed versus non-blood-fed. Differences for enzyme expression were significant for blood-feeding (F = 2.72, P = 0.046) and time (F = 5.37, P = 0.002) **B.** Fold change of microarray expression data (GEO2R GSE22339(Dissanayake et al., 2010)) for tRNA-modifying enzymes. 30 tRNA-modifying enzyme transcripts were evaluated post blood-feeding in *A. aegypti*. Enzymes significantly affected PBF are indicated by * (linear model, P-value < 0.05, outputs listed in Supp Table 4).

Purified tRNA (3μg per sample) was digested using a series of nucleases as previously reported (Kelley et al., 2022a, 2022b). The resulting nucleosides were dried and resuspended in mobile phase A. An internal standard (2-bromo-deoxycytidine, *m/z* = 306.0078) was added to each sample to ensure uniformity of matrix effects.

Mobile phase A (MPA) was composed of 5.3 mM ammonium acetate (pH = 4.5), and mobile phase B (MPB) consisted of acetonitrile/water (40:60) with 5.3 mM ammonium acetate (pH = 4.5). Separation of nucleosides was performed via reverse-phase liquid chromatography with a high strength silica column (Acquity UPLC HSS T3, 1.8 μm, 1.0 mm x 100 mm, Waters) using an ultra-high-performance liquid chromatography system (Vanquish Flex Quaternary, Thermo Fisher Scientific). The flow rate was 100 μL min^-1,^ and the column temperature was 30°C. An Orbitrap Fusion Lumos Tribid mass spectrometer (Thermo Scientific) with a heated-electrospray ionization (H-ESI) source was used for data acquisition. The analyses were carried out in positive mode and a full scan with the *m/z* range of 220 - 900. The LC-MS/MS method and parameters were the same as previously reported (Kelley et al., 2022b).

Data processing was performed using Qual Browser in Xcalibur 3.0 (Thermo Scientific). Modifications were identified by retention time, precursor ion *m/z*, and fragment ion *m/z*. To account for potential variations in tRNA amounts, the nucleoside extracted ion chromatogram (XIC) peak areas were normalized with the summation of the canonical nucleoside (A, G, C, U) peak areas to generate a normalized abundance. Three biological replicates were considered for each time point. For heatmaps, the fold change for each nucleoside was calculated using the normalized peak areas, and NBF levels were considered the initial measurement. A significant difference in levels of a given nucleoside was determined using the normalized peak areas across time points by a general linear model (lme4 package in R) with an output P-value < 0.05 (**Supp Table 3**) and subjected to further statistical testing outlined in the following sections.

### Oligonucleotide analysis using LC-MS/MS to confirm Tyr anticodon modification state

tRNA (5μg) isolated from 12 hours post-blood feeding (PBF) was digested into oligonucleotides for modification mapping experiments. This time point was selected as it aligns with the upregulation of vitellogenin mRNA and Tyr tRNA transcripts (Isoe and Hagedorn, 2007). Two nucleases, RNase T1 and cusativin, were used to improve sequence coverage and confidence in the MS data interpretation (Addepalli et al., 2017; Ross et al., 2017). Ribonuclease T1 cleaves at the 3’- terminus of all guanosine residues, and cusativin preferentially cleaves at the 3’- terminus of all cytidine residues. Both enzymes generally do not cleave at modifications, and cusativin does not cleave consecutive Cs (Addepalli et al., 2017). Because the enzyme selectivity is known, the predicted MS digestion products can be targeted during subsequent LC-MS/MS analysis.

RNase T1 was obtained commercially (Worthington Biochemical, Lakewood, NJ), and recombinant cusativin was overexpressed and isolated in-house (Addepalli et al., 2017). Digestion conditions consisted of 120 mM ammonium acetate and enzyme. RNase T1 (50U/2μg tRNA) and tRNA were incubated at 37°C for 120 min. Cusativin (1μg enzyme/2μg tRNA) and tRNA were incubated at 62°C for 90 min. After digestion, oligonucleotides were dried and resuspended in MPA.

The oligonucleotides were separated using an Xbridge BEH Amide column (2.1 x 150mm, 2.5um, Waters) with an Ultimate 3000 UHPLC (Thermo Scientific) coupled to an LTQ-XL mass spectrometer (Thermo Scientific). The chromatography gradient was performed at 220 μL min-1 flow rate with MPA (70/20/10 v/v H_2_O/acetonitrile/methanol) and MPB (30/60/10 v/v H_2_O/acetonitrile/methanol) with 50 mM ammonium acetate in both mobile phases. The following gradient was used: 5% MPA (0 - 2.5 min), 70% MPA (2.5 - 32.5 min), and then re-equilibration (32.5 - 52.5 min). The resolved digestion products were detected in negative ion mode through ESI conditions of capillary temperature 275 °C, spray voltage of 3.7 kV, and 35, 14, and 10 arbitrary flow units of sheath, auxiliary, and sweep gas, respectively. The initial scan event contains a full scan from *m/z* 600 to 2000, followed by four data-dependent scans triggered by the most abundant precursors from the initial scan event. The *m/z* values of the RNase T1 and cusativin digestion products and fragment ions for the modified and unmodified Tyr tRNA anticodon were predicted using the Mongo Oligo mass calculator (Rozenski, 1999). Mass spectral data was analyzed by manually evaluating each MS/MS spectrum, and oligonucleotide samples were run in duplicate.

### Assessing mature and pretRNA transcript abundance using small RNAseq

Data from small RNA sequencing in the fat body of *A. aegypti* was reprocessed to target mature tRNA and pretRNA transcripts (NCBI Bioproject: PRJNA685230). The tRNA gene list was identified using tRNA-scan 2.0 on the *A. aegypti* genome (Chan et al., 2021; Giraldo-Calderón et al., 2015). The candidate tRNAs were passed through the recommended high confidence eukaryote filter to finalize the gene list (Chan et al., 2021). Isodecoders were manually identified and exact copies of genes were removed before pretRNA mapping. There were 55 and 158 transcripts for mature tRNA and pretRNA, respectively. Three tRNA isoacceptors contained introns: Ile-TAT, Leu-CAA, and Tyr-GTA. For mature tRNA analysis, sequences were generated by the removal of introns and adding CCA to the 3’ end (Hou, 2010; Yoshihisa, 2014). Reads were trimmed using Trimmomatic (version 0.38.0) at default settings (Bolger et al., 2014; Jalili et al., 2020). The trimmed reads were aligned to mature tRNA and pretRNA sequences using Sailfish (version 0.10.1.1) under suggested settings to generate transcripts per million (TPM) (Bolger et al., 2014; Patro et al., 2014). Expression of a tRNA gene was defined as having a TPM > 1. A transcript with TPM < 1 was considered not expressed. Differential expression of tRNAs was determined using DEseq2 (version 2.11.40.7) with an FDR of 0.05 (Jalili et al., 2020; Love et al., 2014).

### Isolation and detection of modifications in the fat body by LC-MS/MS

Fat bodies were collected from mosquitoes by separating the head and thorax from the abdomen, removing internal organs (i.e., Malpighian tubules, ovaries, guts), and placing the pelt (fat body plus a small amount of cuticle) in RNAlater (Hagedorn, 1974; Romoli et al., 2021). Approximately 60-80 pelts were used for each biological replicate, and three biological replicates were collected at each time point. The pelt contains the abdominal cuticle and fat body, which we defined as “fat body” as this is a majority of the biological active tissue (Gulia-Nuss et al., 2011; Price et al., 2011; Romoli et al., 2021; Valzania et al., 2018). The time points considered were NBF, 12H, 24H, 48H, and 72H, which were fed as previously described. Pelts were transferred to bead beater tubes, and the Tri reagent protocols were used to isolate total RNA (Kelley et al., 2022b). LiCl precipitation separated tRNA by resuspending the dried total RNA pellet in 100 μL of 4.4M LiCl. The mixture was incubated at -20°C for 1 hour and then centrifuged for 15 min at 15,000 rpm at room temperature. The supernatant was collected into a new tube, and 0.5X volume of 7.5M ammonium acetate and 3X volume of 100% ethanol was added. After mixing, the samples were stored overnight at -20°C. The next day, the tRNA was precipitated by centrifuging for 30 minutes at 12,000 rpm at 4°C. The pellet was washed with 75% ethanol, centrifuged (15 minutes, 12,000 rpm 4 °C), and dried on the bench for 30 minutes before assessing concentration via Nanodrop (Thermo Scientific).

Purified tRNA samples were digested as described for the whole-body analyses and resuspended in mobile phase A with the aforementioned internal standard. The same LC method and column were used as described in the whole-body modification detection. Data acquisition occurred using a Thermo Scientific TSQ Quantiva Triple Quadrupole mass spectrometer. Selected reaction monitoring (SRM) was performed for all of the modifications identified previously (Kelley et al., 2022b) using predetermined SRM transitions (**Supp Table 2**). Analysis was done using an H-ESI source in positive polarity, with a Q1 and Q3 resolution of 0.7, and Dwell Time of 300 ms. For collision induced dissociation (CID) fragmentation, nitrogen gas was used at 1.5 mTorr; sheath gas, auxiliary gas, and sweep gas of 45, 4.5, and 1.2 arbitrary units, respectively; ion transfer tube temperature of 267 °C; vaporizer temperature of 180 °C; and spray voltage of 3.5 kV.

### Statistics

Data processing for LC-MS/MS was conducted in the same manner for fat body and whole-body samples, and significant differences in modification levels and tRNA modifying enzyme expression levels were determined using a linear regression model (lm function in the lme4 R package (Bates et al., 2015)) and in Supp Table 4 and 5. Expression values for tRNA-modifying enzymes and tRNA modification levels in relation were examined for significant interactions between time and treatment with a multivariate multiple regression model where seventeen modifications and thirty enzymes were evaluated (Kelley et al., 2022b). The data was considered in combinations of modification and single enzyme transcript expression levels for modifications with multiple potential enzymes. An interaction was considered significant when the P-value < 0.05 **Supp Table 3**.

Non-metric multidimensional scaling (NMDS) statistics were performed using anosim (Analysis of Similarity, function “anosim” in R in “vegan” package, 999 permutations) with Bray-Curtis distribution to determine the difference between blood-feeding status, time post blood-feeding, and relationship for modifications and enzymes. Enzyme transcript abundance and modification levels were subjected to permanova (Permutational Analysis of Variance, function “adonis” in R in “vegan” package, 999 permutations) to test differences in transcript levels and tRNA modification abundance between blood-feeding status and time post blood-feeding with Euclidean method. A Fisher’s exact test was used on the percentage of codon frequency to confirm the previously reported enrichment of Tyr codons in vitellogenin (Isoe and Hagedorn, 2007) and compared to reference transcripts (ribosomal proteins). Codon frequency measurements were determined from selected groups of transcripts (i.e., ribosomal protein, vitellogenin) in the Codon Usage Database (Nakamura et al., 2000). Enrichment of tyrosine codon frequency was compared to alanine codon frequency and a P-value < 0.05 was considered significant.

## Results

### Transcript levels of tRNA-modifying enzymes are impacted by blood meal and vitellogenesis

Thirty tRNA-modifying enzyme transcripts were identified and quantified for differences in expression levels following a blood meal (Kelley et al., 2022b). Non-metric multidimensional scaling (NMDS) revealed that enzyme expression patterns were temporally significant following a blood meal for blood meal status (F-stat = 2.72, P-value = 0.046) and time (F-stat = 5.37, P-value = 0.002) (**Figure 1A**). 72H post blood-feeding samples show the closest relationship to non-blood-fed samples in NMDS. This indicates the response to blood-feeding is different in the early times following a blood meal compared to later times when the blood meal is digested.

Of the thirty enzymes considered, transcript levels of fifteen enzymes were significantly different post blood-feeding (**Figure 1B**), where the majority of differentially expressed transcripts were methyltransferases (60%). The other enzymes with transcriptional differences include pseudouridine synthase 1 (PUS1) and PUS7, which are responsible for the modification of pseudouridine (Ψ) and dihydrouridine synthase 3L (DUS3L) and DUS2, which catalyze the formation of dihydrouridine (D). Two anticodon modifiers were also differentially expressed: TP53RK and QTRT2, which correspond to *N6*-threonylcarbamoyladenosine (t^6^A) and queuosine (Q), respectively.

### Anticodon modifications are affected post blood meal consumption

As tRNA-modifying enzyme transcripts are impacted, we examined if tRNA modifications also vary following a blood meal. Mosquito tRNA modifications identified here are the same as those previously reported in female mosquitoes (Kelley et al., 2022b) except for mcm^5^s^2^U, which was too low of abundance to be consistently detected. NMDS revealed that blood-feeding status primarily affects modification levels (**Figure 2A**). The relationship of modification levels with blood-feeding is significant (F-stat = 2.72, P-value = 0.046), which suggests blood meal consumption generally shifts tRNA modification patterns. In contrast, modification levels with respect to time were not significant. However, when the 72H time point is removed from statistical consideration, modification patterns are also substantial for time (F-stat = 2.977, P-value = 0.035).

**Figure 2:**
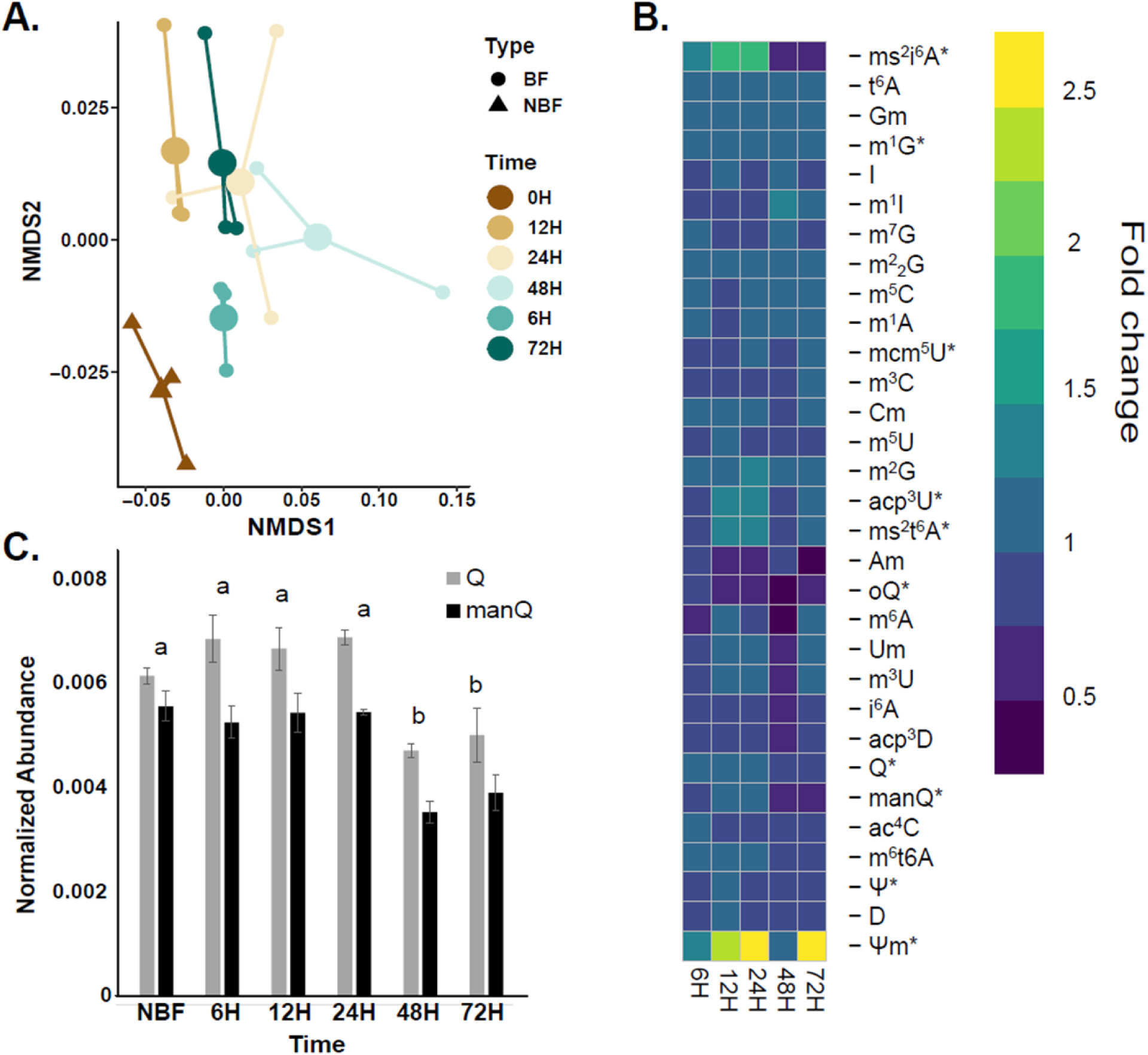
tRNA modification levels are altered post blood meal. **A.** NMDS of tRNA modification abundances, which demonstrates distinct differences based on blood meal status. Differences in modification levels were significant for blood-feeding (F = 3.39, P = 0.035) but not significant for time. **B.** Heatmap of the fold change of relative abundances of tRNA modifications post blood-feeding. Significance was determined by a linear model and is denoted by * (P-value < 0.05). **C.** The normalized abundance of Q and manQ levels across the blood-feeding time course. The peak areas of Q and manQ were normalized with the summation of the canonical peak areas. Significance was determined by a linear model (P-value < 0.05), and post-hoc differences were determined showing changes only at 48 and 72H.

Many modifications remain at similar levels to control for two days following a blood meal (**Figure 2B**). Altogether, twelve modifications are affected post blood meal, which include: Ψ, Ψm, 1-methylguanosine (m^1^G), 5-methoxycarbonylmethyluridine (mcm^5^U), *N6*-isopentenyladenosine (i^6^A), Q, manQ, epoxyqueuosine (oQ), ms^2^i^6^A, 3-(3-amino-3-carboxypropyl)uridine (acp^3^U), t^6^A, and ms^2^t^6^A. For some modifications, such as Q and manQ, levels are reduced around 48H post blood meal and remain low at 72H compared to non-blood-fed levels (**Figure 2C**). The other modifications that demonstrate this pattern include 2’-O-methylpseudouridine (Ψm), ms^2^i^6^A, and ms^2^t^6^A.

Modification abundances and enzyme expression patterns significantly interacted with time or treatment for ten modifications (**Table 1**). These modifications were Ψ, D, m^3^C, Cm, m^1^A, m^2^G, Gm, m^1^G, Q, and t^6^A. Intriguingly, six of the ten significant modifications are methylations, and the methyltransferases associated with these modifications were also differentially expressed.

### Tyrosine codon enrichment in vitellogenin and modification with manQ

While several modifications demonstrated altered levels post-blood meal, anticodon modification changes were of primary interest due to potential impacts on decoding. Two anticodon modifications that varied post blood meal were Q and manQ. These modifications are typically found on tRNAs of four amino acids (aspartic acid, histidine, tyrosine, and asparagine) (Boccaletto et al., 2018; Hillmeier et al., 2021). As tRNA modifications are critical to decoding codons, we examined the codon composition of vitellogenin transcripts in *A. aegypti*. Tyrosine codon frequency was doubled in vitellogenin transcripts compared to references of ribosomal protein transcripts and codon usage in the whole genome (**Figure 3A**). Overall, the preferences for specific tyrosine codons (i.e., UAC or UAU) are similar in the genome, ribosomal proteins, and vitellogenin genes (**Supp Figure 2**).

**Figure 3:**
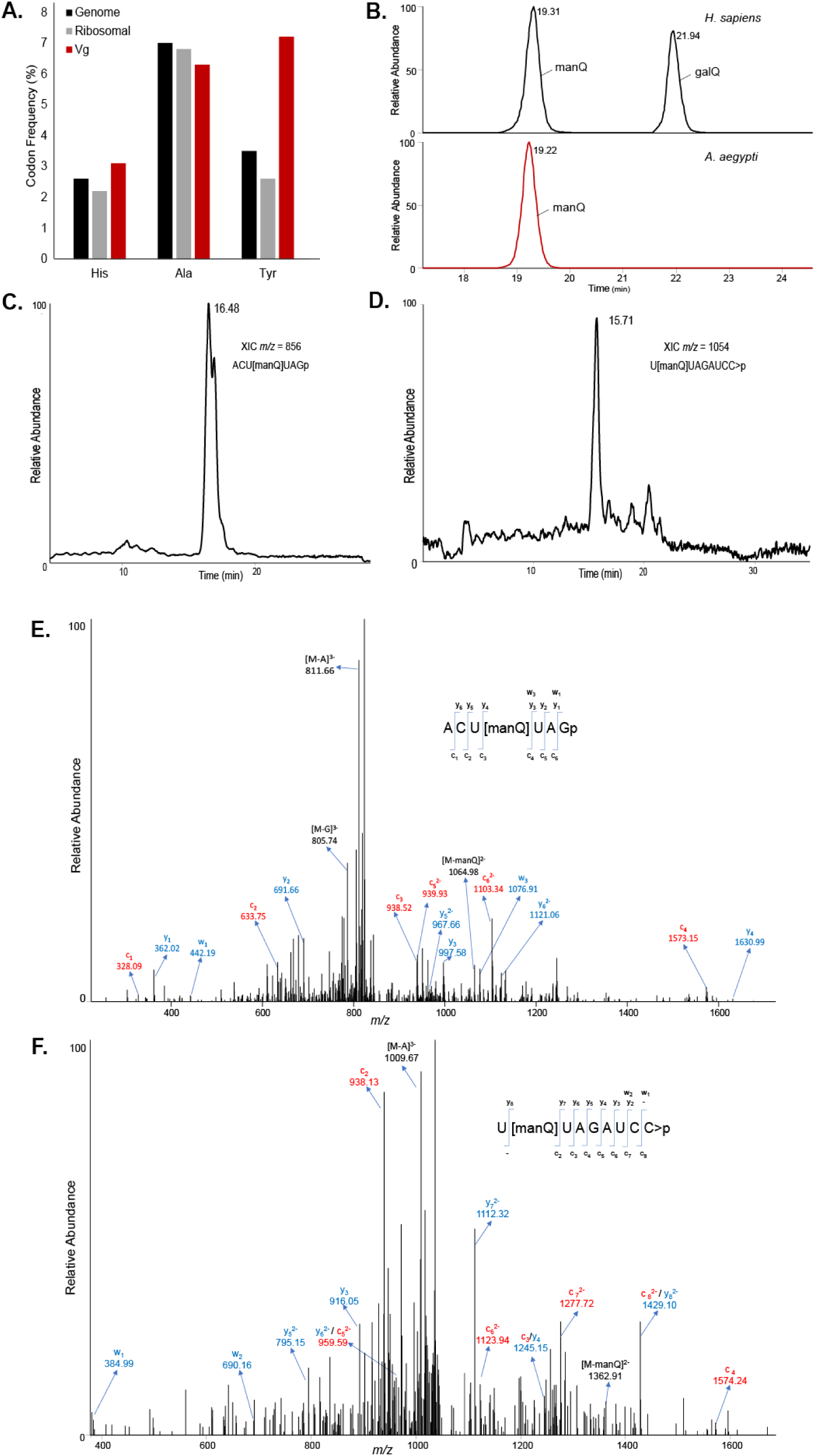
Mannosyl-queuosine (manQ) is the wobble position modification on Tyr tRNA in *A. aegypti*. **A.** Enrichment of tyrosine codon usage in vitellogenin indicates Tyr tRNA is likely important for meeting the codon demand. A Fisher’s test showed enrichment of Tyr codons in Vg transcripts compared to ribosomal protein transcripts as a reference (P-value < 0.01). **B.** XIC of *m/z* = 572.2204 top panel demonstrates that galQ and manQ are present in human tRNA. The isomers are typically detected together; however, only manQ is detected in *A. aegypti (Kelley et al., 2022b)*. **C.** XIC of T1 generated oligonucleotide ACU[manQ]UAGp with *m/z* = 856from the anticodon loop of Tyr tRNA. **D.** XIC of the cusativin oligonucleotide U[manQ]UAGAUCC>p with m/z = 1054. Both digestion products indicate that manQ is at position 34 of Tyr tRNA in *A. aegypti*. **E.** Mass spectrum for the oligonucleotide ACU[manQ]UAGp with the labeled fragment ions. **F.** Mass spectrum for the oligonucleotide U[manQ]UAGAUCC>p with the associated fragment ions.

The anticodon wobble position modification on tyrosine tRNA is typically Q or galactosyl-queuosine (galQ) (Boccaletto et al., 2018; Okada and Nishimura, 1977); however, the modification galQ is not present in *A. aegypti* tRNA (**Figure 3B**). As galQ has been previously mapped to the anticodon of Tyr tRNA (Boccaletto et al., 2018), it was unclear if Q or manQ were on the anticodon in *A. aegypti*. To determine the anticodon modifications, tRNA was enzymatically digested into predictable oligonucleotides. It is possible to identify and map RNA modifications onto the sequence using the known tRNA sequence, modifications present, and LC-MS/MS (Jora et al., 2019; Ross et al., 2017). The digested oligonucleotides were separated chromatographically and then detected with the mass spectrometer. The oligonucleotide ACU[manQ]UAGp (*m/z* = 856) was observed with a retention time of approximately 16.5 minutes (**Figure 3C**). Likewise, the cusativin digestion product U[manQ]UAGAUCC>p (*m/z* = 1054) demonstrated a retention time of 15.7 minutes (**Figure 3D**).

After chromatographic separation, the oligonucleotide is fragmented by CID, and the pattern of fragmentation for oligonucleotides is well-established (Gaston and Limbach, 2014; Kowalak et al., 1993; Ross et al., 2017). The c- and y-type fragment ions are determined by the sequential dissociation of the oligonucleotide phosphodiester backbone. With this approach, the modification manQ was mapped to position 34 in Tyr tRNA by identification of the oligonucleotide ACU[manQ]UAGp (*m/z* = 856) (**Figure 3E**). In agreement, the Tyr anticodon was detected in cusativin digestion with the oligonucleotide U[manQ]UAGAUCC>p (*m/z* = 1054) (**Figure 3F**). The digestion products are unique to only Tyr tRNA sequences, improving confidence in the identification of the Tyr anticodon (Wetzel and Limbach, 2012). While m^1^G has been reported at position 37 of Tyr tRNA in other organisms (Boccaletto et al., 2018), no digest products containing m^1^G37 were detected for the Tyr anticodon sequences. If manQ was not present, an unmodified G was identified, and there were no digestion products containing oQ or Q from whole-body analyses (**Supp Figure 3**).

### Fat body modifications and mature tRNA transcripts are altered post blood-feeding

Vitellogenin is synthesized in the fat body; as such, there is likely a higher demand for Tyr tRNA than other tissues following a blood meal due to Tyr codon frequency. Only five of 55 mature tRNAs examined were differentially expressed in the fat body in all time points post blood-feeding. Two tRNAs, Ile-AAT-1 and Thr-CGT-1, were decreased across time points. The other three differentially expressed mature tRNAs, Tyr-GTA-1, Ala-AGC-3, and Ala-TGC-1, were consistently elevated in response to blood-feeding. At 6H PBF, tRNA Lys-TTT-1 is heightened but returns to non-blood-fed levels by 12H PBF. Notably, the transcript levels of Tyr tRNA in the fat body are elevated and remain higher PBF (**Figure 4A**), correlating with the enrichment of Tyr codons in vitellogenin transcripts. While Ala-AGC-3-1 followed a similar trend to Tyr-GTA-1-1, there is no enrichment of Ala codons in vitellogenin, suggesting Ala tRNA could be involved in other biological processes occurring in the fat body PBF.

**Figure 4.**
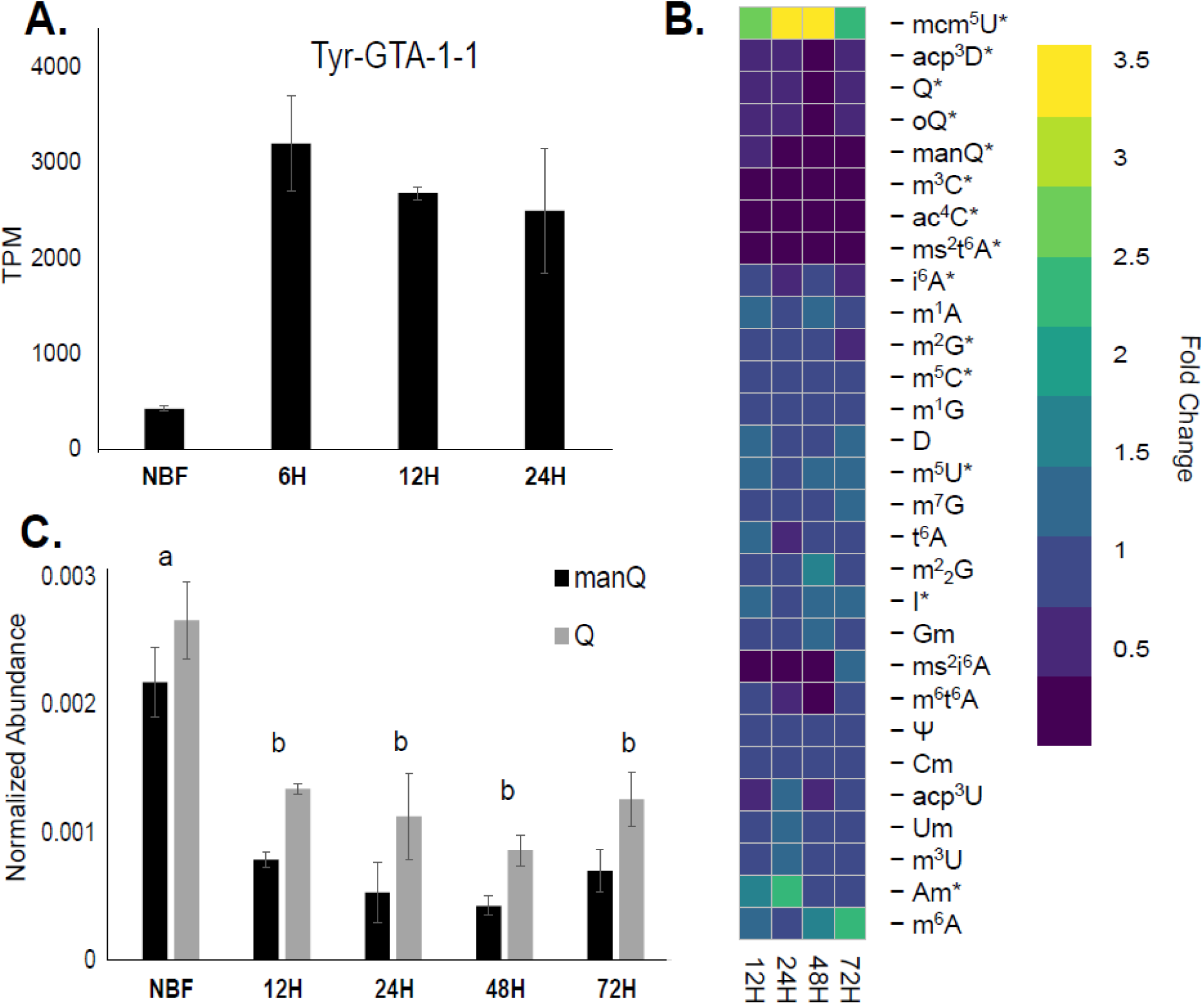
Tyrosine mature tRNA transcripts are in high abundance but anticodon modification level is lowered post blood-feeding in the fat body. **A.** Tyr mature tRNA transcripts are five-fold higher post blood-feeding in the fat body. Significance determined using DEseq2, FDR < 0.05. **B.** The wobble position modifications, Q and manQ, are significantly lower post blood-feeding in the fat body. Significance determined by linear model (P-value < 0.05). **C.** Heatmap of the fold change of relative abundances of tRNA modifications post blood-feeding in the fat body. Significance was determined by a linear model and is denoted by * (P-value < 0.05)

As our oligonucleotide analyses indicate manQ is the Tyr tRNA anticodon modification, we performed tRNA modification analysis of the fat body at time points post blood-feeding that correlate with critical times for vitellogenesis (Isoe and Hagedorn, 2007). Two modifications in whole body samples were not consistently detectable in the fat body: Ψm and 1-methylinosine (m^1^I). In total, 14 modifications exhibit altered levels in the fat body post blood-feeding, including oQ, Q, manQ, acp^3^D, ms^2^t^6^A, m^3^C, ac^4^C, mcm^5^U, m^5^U, I, Am, m^2^G, i^6^A, and m^5^C (**Figure 4B**). Modifications in higher abundances PBF were mcm^5^U, m^5^U, I, and Am. Intriguingly, mcm^5^U has been mapped to the wobble position on Lys-TTT tRNAs (Fernández-Vázquez et al., 2013; Huang et al., 2005), which is one of the tRNAs elevated in the fat body PBF. mcm^5^U has been implicated in stress responses in other organisms due to its roles in translational efficiency and fidelity (Begley et al., 2007); however, the biological function of this modification and others elevated in the fat body is unclear. Of particular interest, levels of manQ and Q were dramatically decreased in relative abundance post blood-feeding in the fat body (**Figure 4C**), and this occurred in several other modifications (oQ, acp^3^D, ms^2^t^6^A, m^3^C, and ac^4^C) as well. Thus, there is an increase in Tyr tRNA in the fat body but no corresponding increase in manQ levels, suggesting under modified tRNAs PBF.

### Differential expression of Tyr pretRNA post blood-feeding

Regardless of tRNA modification status, the measurement of tRNA modification abundance is relative to the entire tRNA pool. Therefore, we investigated whether pretRNA levels differed in non-blood fed and post blood-feeding. When RNA polymerase III first transcribes tRNA, the transcript contains leader, trailer, and, occasionally, intron sequences. These sequences are removed enzymatically, and the transcript is further processed to form the mature tRNA for translation. The presence of an intron allows for distinguishing the tRNA genes that are recently transcribed (pretRNA) compared to the mature tRNA pools.

Tyr tRNA genes canonically contain an intron, allowing small RNAseq data to be analyzed following blood-feeding to examine changes in Tyr pretRNA levels. In *A. aegypti*, two other tRNAs contain introns, Ile-TAT and Leu-CAA, which were considered in the analyses for comparison to Tyr pretRNA changes. There was a consistent elevation of Tyr pretRNA post blood-feeding with eight Tyr pretRNA only expressed following a blood meal (**Figure 5A**). Seven pretRNAs were differentially expressed following a blood meal with the 12H time point having six Tyr pretRNA with enriched expression (**Figure 5B**). Three of the thirty Tyr pretRNAs were consistently elevated post blood-feeding: Tyr-GTA-15-1, Tyr-GTA-9-2, and Tyr-GTA-1-4. (**Figure 5C**). Expression levels of other pretRNAs considered, Ile-TAT and Leu-CAA, were not significantly impacted by blood meal status. These results suggest that Tyr tRNA expression increases in the fat body following a blood meal.

**Figure 5.**
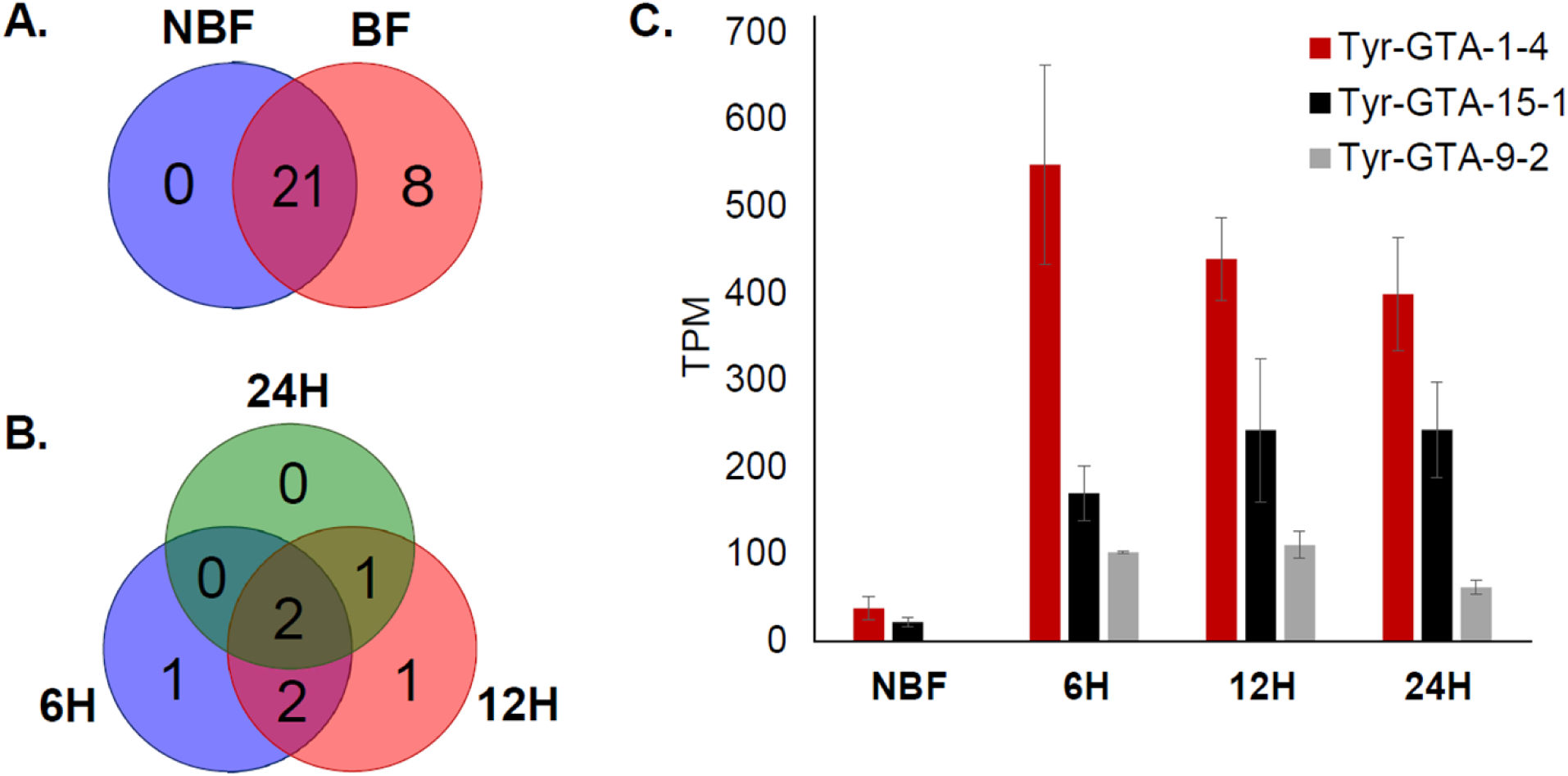
Tyrosine pretRNA are upregulated post blood-feeding in the fat body. **A.** Nine of thirty Tyr pretRNA are expressed exclusively post blood-feeding. Expression was defined by having a TPM > 1. A transcript with TPM < 1 was considered not expressed. **B.** Tyr pretRNA differential expression is dependent on the hours post blood-feeding. Significance determined by DEseq2 FDR < 0.05 **C.** Three Tyr pretRNA were consistently higher across blood-feeding time points. Significance determined by DEseq2 FDR < 0.05.

## Discussion

Numerous physiological and molecular changes, such as transcriptome and proteome remodeling, occur in mosquitoes in response to blood feeding (Camargo et al., 2020; Hixson et al., 2022; Matthews et al., 2016; Sanders et al., 2003). Before blood meal ingestion, a pre-vitellogenic period of arrest occurs and is hormonally regulated (Fallon et al., 1974; Wu et al., 2020). Immediately following blood meal consumption, a complex interplay of molecular reactions begins, including the induction stress responses (Benoit et al., 2019, 2011; Sterkel et al., 2017), activation of the target of rapamycin (TOR) pathway (Attardo et al., 2005; Hansen et al., 2014), and synthesis of egg yolk proteins such as vitellogenin (Hansen et al., 2014). Along with the characterized responses, we have identified that alterations in tRNA expression and modification occur post blood-feeding and during vitellogenesis in *A. aegypti*.

There is a global response to blood-feeding in tRNA modification levels and tRNA-modifying enzyme transcripts. Some modifications affected by a blood meal support a heat shock response. Methylations (i.e., m^1^G, Ψm) and acp^3^U are implicated in the stability of tRNA at higher temperatures (Abou Assi et al., 2020; Hori et al., 2018; Takakura et al., 2019). Methyltransferase enzymes are differentially expressed with transcript levels for several enzymes (i.e. TRMT61A, TRMT5, TRMT10C) increasing following the blood meal. Furthermore, there are changes to modifications located on the anticodon loop, particularly at the wobble position and the position adjacent to the anticodon or positions 34 and 37, respectively. For example, the position 37 modifications t^6^A and ms^2^t^6^A are consistently elevated PBF and have critical roles in improving codon-anticodon interactions, preventing frameshifting, and improving translational fidelity (Murphy et al., 2004; Perrochia et al., 2013; Stuart et al., 2000). The expression levels of an enzyme involved in t^6^A modification are lowered PBF, but it is possible the pre-blood-feeding levels of the enzyme are sufficient for modification. Wobble modifications affect the decoding of mRNA codons and, thus, the accuracy and speed of translation (Agris et al., 2007). The mosquito’s wobble position modifications impacted by blood-feeding were mcm^5^U, oQ, Q, and manQ. Levels of mcm^5^U fluctuate randomly post blood-feeding, possibly due to shifts in the whole body tRNA pool composition (Rak et al., 2021). In the case of Q and manQ, the modification levels are steady and decrease at 48H and 72H PBF, which coincides with the termination of vitellogenesis and oviposition of eggs (Hansen et al., 2014; Isoe and Hagedorn, 2007; Zhu et al., 2003). Overall, the shifts in modifications emphasize the general importance of translational fidelity and efficiency as well as the stability of tRNA in hours following blood-feeding in mosquitoes.

Many accounts of tRNA modifications being remodeled to meet codon demand of transcripts have been noted during specific conditions (Chionh et al., 2016; Dedon and Begley, 2014; Endres et al., 2015; Huber et al., 2019). The documented enrichment of tyrosine codons in vitellogenin (Isoe and Hagedorn, 2007) demonstrated there may be a demand for tRNA modification, particularly in the fat body where vitellogenesis occurs (Bryant and Raikhel, 2011). Typically, galQ is the wobble position modification on Tyr tRNA in eukaryotes (Boccaletto et al., 2018; Okada and Nishimura, 1977); however, this modification was not present in *A. aegypti* tRNA (Kelley et al., 2022b). Instead, we identified manQ as the wobble position modification on Tyr tRNA and hypothesized levels of manQ would be elevated in the fat body PBF. Subsequent modification analysis revealed that relative levels of manQ and Q are significantly decreased in the fat body PBF. Similarly, a subset of other modifications diminishes in fat body tRNA following blood-feeding. Increases in four modifications are also observed, but the biological implications for these modifications are unclear. Ultimately, manQ relative abundance was consistently lower during time points coinciding with vitellogenin production in the fat body. This suggests an alternative process is required to account for tyrosine codon demand during vitellogenin production.

The abundance of tRNA transcripts is another strategy to meet codon demand in response-specific transcripts (Aharon-Hefetz et al., 2020; Torrent et al., 2018). In mosquitoes, mature Tyr tRNA transcripts were consistently higher post blood meal and correlated with times of heightened vitellogenin expression (Isoe and Hagedorn, 2007). With the elevation of Tyr tRNA but the relative decrease of manQ modification, there is likely an increase in tRNA without manQ at the wobble position. In human cells, loss of wobble position modification phenotypes can be compensated by increased expression of the associated tRNA (Bauer and Hermand, 2012). Lower levels of Q in tRNA of human cells lead to protein aggregation and activation of the unfolded protein response due to ribosome stalling at Q-decoded codons (Tuorto et al., 2018). It is still being determined if lack of manQ could impact the quality of vitellogenin produced during protein synthesis. Furthermore, the availability of Q as a micronutrient may be altered by consuming a blood meal. In eukaryotes, Q must be salvaged from diet or the bacteria in the microbiome (Zallot et al., 2014), which suggests the availability of this micronutrient could influence tRNA modification in mosquitoes. All in all, functional studies are necessary to elucidate the role Q has on vitellogenesis in mosquitoes.

Tyr tRNA genes contain an intron sequence present in newly transcribed precursor Tyr pretRNA (Chan and Lowe, 2016; van Tol and Beier, 1988). Utilizing this feature, we assessed Tyr pretRNA levels, which were elevated post blood-feeding. Interestingly, eight of the twenty-nine Tyr pretRNAs were exclusively expressed post blood-feeding in the fat body. In addition, several Tyr pretRNAs are increased at specific time points post blood-feeding. Five Tyr pretRNAs are immediately increased 6H PBF, which indicates some Tyr tRNA genes are transcribed during the initial stages of vitellogenesis. After 12H, six Tyr pretRNAs are transcribed and three pretRNAs remain at significantly heightened levels by 24H PBF, further demonstrating temporal Tyr tRNA gene transcription. Intriguingly, two of the tRNAs expressed after blood-feeding, Tyr-GTA-10-1 and Tyr-GTA-16-1, have unique sequences 3’ of the intron, and no other Tyr tRNA contains them. Notably, sequence differences in isodecoders often occur at locations of RNA polymerase III contacts to initiate gene expression, which likely contributes to tissue-specific regulation of tRNA gene expression (Pinkard et al., 2020). Further investigation on the tRNA expression of other tissues is necessary to determine if isodecoder expression is tissue-specific and dependent on blood-feeding in mosquitoes. Overall, tRNA gene transcription is likely promoted by blood meal consumption and occurs in an isodecoder specific manner to synthesize vitellogenin.

Activation of the evolutionarily-conserved TOR signaling cascade in the fat body promotes protein synthesis and vitellogenesis (Attardo et al., 2005; Brandon et al., 2008; Hansen et al., 2014). In other eukaryotes, induction of TOR signaling inhibits the repressor for RNA polymerase III (Marshall et al., 2012; Upadhya et al., 2002), which is responsible for the transcription of tRNA genes (Leśniewska and Boguta, 2017; Turowski and Tollervey, 2016; Wei et al., 2009). Therefore, activation of TOR likely promotes the transcription of tRNAs and other protein synthesis components. In *Drosophila*, the TOR pathway directly impacts tRNA synthesis, which is controlled by nutrients and hormones, particularly in the fat body (Marshall et al., 2012). Furthermore, the transcription of tRNA genes is a conserved process that depends on nutrient and stress signaling (Moir and Willis, 2013). The amino acid influx brought on by the blood meal is critical to induction of TOR signaling (Attardo et al., 2005; Hansen et al., 2004), elevates the global rate of protein synthesis (Thomas and Hall, 1997; Wolfson and Sabatini, 2017), and likely promotes tRNA transcription. Ultimately, increased amino acid levels promote the TOR signaling cascade and likely play a role in Tyr tRNA transcription following blood-feeding; however, additional research is required to establish a direct link.

Previous work in mosquitoes suggests that the molecular mechanisms involved in production of vitellogenin are transcriptionally regulated (Fallon et al., 1974; Hagedorn, 1974). Here, we demonstrate codon usage, tRNA modification, and transcripts of tRNA are impacted by the blood-feeding response and subsequent vitellogenesis. Thus, we propose a model where blood consumption induces Tyr tRNA transcription in the fat body to account for vitellogenin codon demand (**Figure 6**). Higher levels of Tyr tRNA are produced rapidly and are likely unmodified at the wobble position, causing the relative abundance of Tyr tRNA wobble modification to lower. Analyses via isotope labeling could be used to confirm that newly transcribed Tyr tRNAs are undermodified (Heiss et al., 2021). Altogether, we show that Tyr tRNA levels are increased in the fat body, providing the molecular machinery to allow for increased synthesis of vitellogenin. These studies suggest the inhibition of Tyr tRNA synthesis could be used as a novel mechanism to suppress vitellogenin synthesis in mosquitoes, which warrants further investigation.

**Figure 6.**
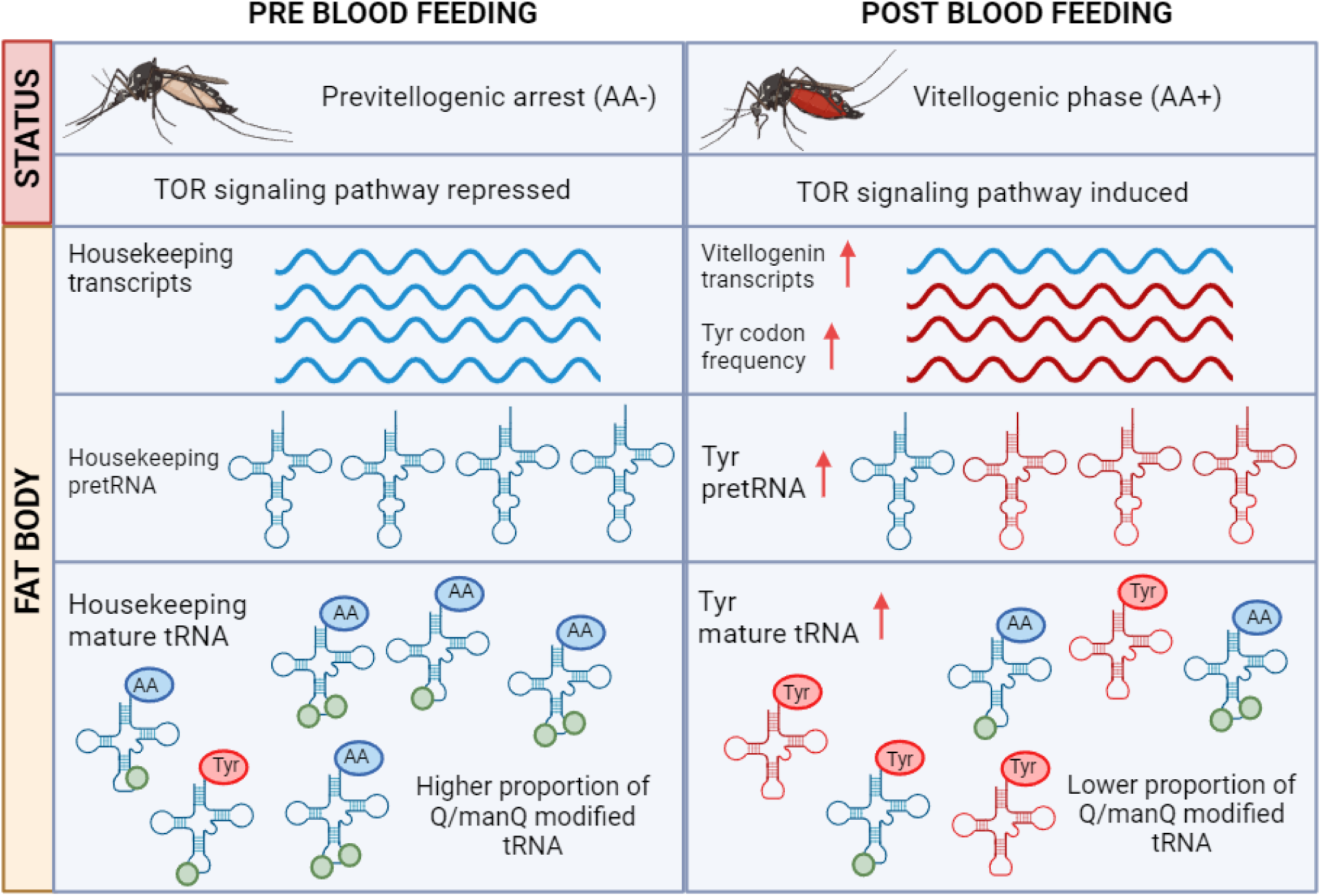
Proposed model of effects of blood-feeding on tRNA in the fat body. Before blood-feeding, the tRNA pool is stable and Q/manQ is modified in the fat body to meet the codon demand of housekeeping transcripts. Post blood-feeding, the TOR signaling pathway is activated and vitellogenin transcripts are increased (Attardo et al., 2005). Enrichment of tyrosine codons in vitellogenin requires a new set of tRNAs. Therefore, pretRNA are remodeled to enrich Tyr tRNA. The increase of tRNA transcripts is also reflected in the mature tRNA. The rapid increase of Tyr tRNA likely results in higher transcripts that are undermodified at the wobble position, which lowers the relative abundance of wobble modifications associated with Tyr tRNA.

## Acknowledgments

Research reported in this publication was partially supported by the National Institute of Allergy and Infectious Diseases of the National Institutes of Health under Award Number R01AI148551 and R21AI166633 (to J.B.B.) and the National Institutes of Health (GM 058843 to P.A.L). Direct support is provided by the National Institute of Allergy and Infectious Diseases of the National Institutes of Health under Award Number R21AI176098. The content is solely the responsibility of the authors and does not necessarily represent the official views of the National Institutes of Health.

