## Supplemental Tables and Figures for "Tyrosine transfer RNA levels and modifications during blood-feeding and vitellogenesis in the mosquito, *Aedes aegypti*"

Supplemental Figures and Tables

**Supplemental Table 1**. Genes of tRNA-modifying enzymes identified in *A. aegypti*. The list of previously identified genes in *A. aegypti* associated with tRNA modification. The microarray IDs from Dissanayake et al 2010 are also listed.

| Gene | Name | Modification | ID_1 | ID_2 |
|---|---|---|---|---|
| AAEL007354 | PUS1 | Ψ | 12230 | 32222 |
| AAEL004663 | PUS3 | Ψ | 19066 | 41711 |
| AAEL003071 | PUS7 | Ψ | 14193 | 17473 |
| AAEL010362 | PUS10 | Ψ | 35068 | 36380 |
| AAEL005898 | DUS2 | D | 671 | 42043 |
| AAEL001171 | DUS3L | D | 5550 | 1.09E+04 |
| AAEL008120 | FTSJ1A | Cm/Gm | 14814 | 30069 |
| AAEL001037 | FTSJ1B | Cm/Gm | 30196 | 31442 |
| AAEL014096 | TRMT44 | Um | 10629 | 31124 |
| AAEL011199 | TRMT13 | Cm/Am | 11230 | 29865 |
| AAEL003922 | TRMT11 | m2G | 5108 | 8488 |
| AAEL012932 | METTL6 | m3C | 10058 | 37114 |
| AAEL007084 | METTL2B | m3C | 22856 | 43453 |
| AAEL009603 | TRMT5 | m1G | 18667 | 22767 |
| AAEL008124 | TRMT61A1 | m1A | 5632 | 19910 |
| AAEL003654 | TRMT61A2 | m1A | 11245 | 32926 |
| AAEL008941 | TRMT10A | m1A | 4156 | 34445 |
| AAEL011538 | TRMT10C | m1G/m1A | 5938 | 11440 |
| AAEL006166 | TRDMT1 | m5C | 17034 | 26850 |
| AAEL012520 | NSUN3 | m5C | 34636 | 39482 |
| AAEL010349 | TRMT1 | m22G | 7195 | 33365 |
| AAEL004625 | DTWD1 | acp3U | 9592 | 30264 |
| AAEL010993 | QTRT1 | Q | 9669 | 17759 |
| AAEL010968 | QTRT2 | Q | 8566 | 18414 |
| AAEL001036 | ELP1 | mcm5U | 7937 | 27473 |

| | | | | |
|---|---|---|---|---|
| AAEL004333 | ELP4 | mcm5U | 10072 | 19052 |
| AAEL004571 | OSGEPL1 | t6A | 22530 | 33617 |
| AAEL006221 | TP53RK | t6A | 2144 | 40735 |
| AAEL007313 | TrmO | m6t6A | 16015 | 32419 |
| AAEL002837 | CDK5RAP1 | ms2i6A | 13433 | 27734 |

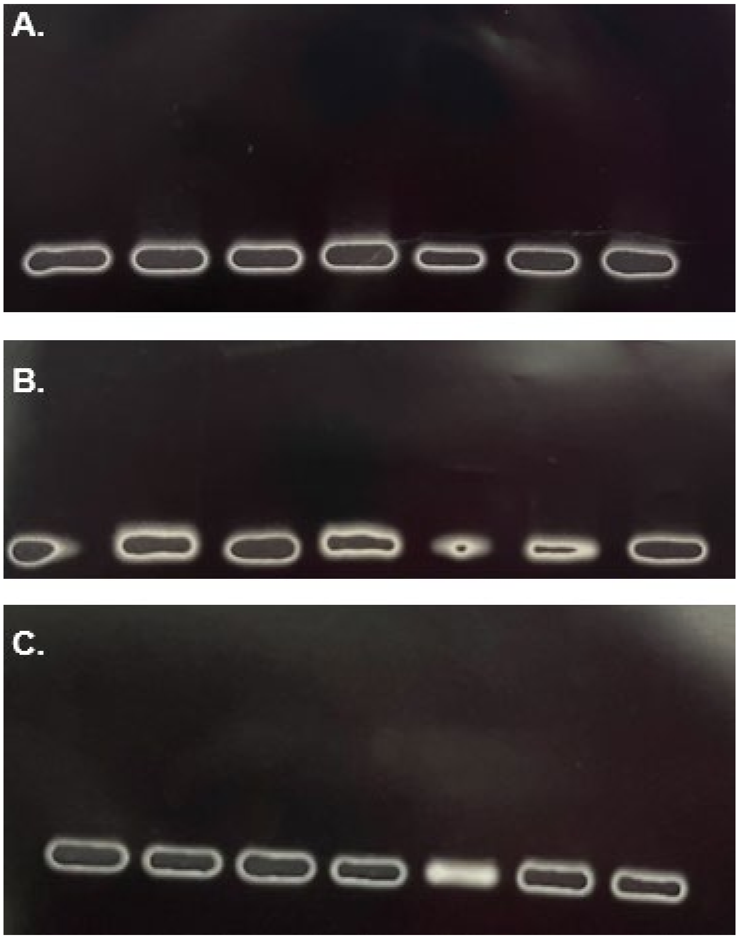


**Supplemental Figure 1. Agarose gel confirming the presence of pure tRNA prior to nucleoside digestion.** In each gel, the first lane on the left is a yeast tRNA standard. A. Order (left to right): standard, NBF_1, NBF_2, NBF_3, 72H_1, 72H_2, 72H_3. B. Order (left to right): standard, 6H_1, 6H_2, 6H_3, 12H_1, 12H_2, 12H_3. C. Order (left to right): standard, 24H_1, 24H_2, 24H_3, 48H_1, 48H_2, 48H_3.

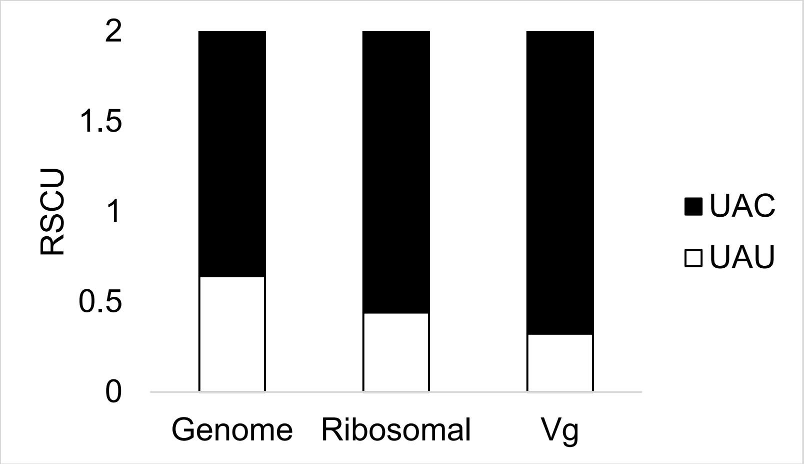


**Supplemental Figure 2. Relative synonymous codon usage (RSCU) of tyrosine codons indicates UAC is the preferred codon.** A RSCU value > 1.5 indicates preference for a codon. A codon is considered unpreferred is the RSCU < 0.5. Despite the same preferred codon, the codon UAU is unpreferred in ribosomal proteins and vitellogenin.

**Supplemental Table 2**. MS parameters for SRM detection of nucleosides in the mosquito fat body.

| Compound | Retention Time (min) | Precursor (m/z) | Product (m/z) | Collision Energy (V) | RF Lens (V) |
|---|---|---|---|---|---|
| Ψ | 2 | 245 | 209 | 10 | 30 |
| C | 3 | 244 | 112 | 12 | 41 |
| D | 3 | 247 | 115 | 10 | 43 |
| m1A | 3 | 282 | 150 | 20 | 74 |
| U | 3.35 | 245 | 113 | 10 | 30 |
| acp3U | 3.71 | 346 | 214 | 10 | 50 |
| acp3D | 2.8 | 348 | 216 | 10 | 50 |
| m3C | 4.07 | 258 | 126 | 14 | 46 |
| m6A | 5.17 | 282 | 150 | 20 | 74 |
| m5C | 5.61 | 258 | 126 | 14 | 46 |
| Ψm | 5.83 | 259 | 223 | 10 | 30 |
| Cm | 7.89 | 258 | 112 | 12 | 41 |
| I | 9.23 | 269 | 137 | 10 | 40 |
| m7G | 9.37 | 298 | 166 | 16 | 50 |
| m5U | 9.55 | 259 | 127 | 10 | 30 |
| G | 11 | 284 | 152 | 14 | 46 |
| m3U | 13.27 | 259 | 127 | 10 | 30 |
| Um | 14.17 | 259 | 113 | 10 | 30 |
| Um | 14.17 | 259 | 127 | 10 | 30 |
| oQ | 15.21 | 426 | 163 | 29 | 84 |
| oQ | 15.21 | 426 | 295 | 12 | 84 |
| Q | 17.6 | 410 | 295 | 16 | 49 |
| Q | 17.6 | 410 | 163 | 31 | 96 |

| | | | | | |
|---|---|---|---|---|---|
| ISTD | 18.7 | 613 | 306 | 12 | 41 |
| Gm | 20.94 | 298 | 152 | 11 | 50 |
| mcm5U | 21 | 317 | 125 | 10 | 30 |
| mcm5U | 21 | 317 | 153 | 10 | 30 |
| mcm5U | 21 | 317 | 185 | 10 | 30 |
| m1I | 21.97 | 283 | 151 | 20 | 74 |
| manQ | 22.94 | 572 | 163 | 35 | 96 |
| manQ | 22.94 | 572 | 295 | 35 | 96 |
| ac4C | 24.16 | 286 | 154 | 10 | 46 |
| A | 24.21 | 268 | 136 | 17 | 63 |
| m1G | 28 | 298 | 166 | 16 | 50 |
| m2G | 30 | 298 | 166 | 16 | 50 |
| m2,2G | 31.2 | 312 | 180 | 15 | 60 |
| m6t6A | 33.25 | 427 | 295 | 31 | 96 |
| Am | 33.39 | 282 | 136 | 17 | 63 |
| mcm5s2U | 34.06 | 333 | 141 | 35 | 96 |
| mcm5s2U | 34.06 | 333 | 169 | 35 | 96 |
| mcm5s2U | 34.06 | 333 | 201 | 35 | 96 |
| ms2t6A | 36.04 | 459 | 208 | 31 | 96 |
| ms2t6A | 36.04 | 459 | 327 | 31 | 96 |
| i6A | 36.54 | 336 | 204 | 31 | 96 |
| i6A | 38.39 | 336 | 148 | 31 | 96 |
| i6A | 38.39 | 336 | 204 | 31 | 96 |
| i6A | 38.39 | 336 | 136 | 31 | 96 |
| ms2i6A | 40.93 | 382 | 250 | 31 | 96 |
| t6A | 34.75 | 413 | 281 | 31 | 96 |

**Supplemental Table 3**. All combinations of interactions between tRNA modification, modifying enzyme expression, time, and blood-feeding. Statistics generated from

| Modification | Gene | Treatment F-statistic | P-value | Time F-statistic | P-value |
|---|---|---|---|---|---|
| Ψ | PUS1 | 70.395 | 5.38E-07 | 6.882 | 0.0001 |
| | PUS3 | 0.87 | 0.445 | 1.8 | 0.125 |
| | PUS7 | 18.03 | 0.0003 | 6.871 | 0.0001 |
| D | DUS2 | 9.57 | 0.003 | 6.1 | 0.0002 |
| | DUS3L | 5.46 | 0.02 | 4.85 | 0.001 |
| m3C | METTL6 | 7.34 | 0.009 | 6.24 | 0.0002 |
| | METTL2B | 1.9 | 0.195 | 5.8 | 0.0003 |
| m5C | TRDMT1 | 1.88 | 0.198 | 1.45 | 0.226 |
| | NSUN3 | 0.233 | 0.795 | 1.77 | 0.131 |
| Cm | FTSJ1A | 0.285 | 0.756 | 2.42 | 0.044 |
| | FTSJ1B | 6.8 | 0.01 | 4.98 | 0.001 |
| | TRMT13 | 0.15 | 0.862 | 2.99 | 0.01 |
| Um | TRMT44 | 0.458 | 0.644 | 2.11 | 0.074 |

| | | | | | |
|---|---|---|---|---|---|
| m1A | TRMT10C | 7.02 | 0.01 | 2.86 | 0.021 |
| | TRMT10A | 0.639 | 0.546 | 2.88 | 0.02 |
| m2G | TRMT11 | 1.63 | 0.239 | 3.14 | 0.01 |
| Gm | FTSJ1A | 1.83 | 0.205 | 1.32 | 0.278 |
| | FTSJ1B | 20.37 | 0.0001 | 2.82 | 0.023 |
| m1G | TRMT10C | 36.96 | 1.31E-05 | 5.32 | 0.0006 |
| | TRMT10A | 21.86 | 0.0001 | 4.97 | 0.001 |
| m22G | TRMT1 | 2.4 | 0.136 | 1.72 | 0.144 |
| mcm5U | ELP1 | 0.816 | 0.467 | 1.92 | 0.102 |
| | ELP4 | 1.01 | 0.395 | 1.94 | 0.1 |
| acp3U | DTWD1 | 0.609 | 0.561 | 2.05 | 0.082 |
| Q | QTRT1 | 0.037 | 0.963 | 1.92 | 0.102 |
| | QTRT2 | 0.379 | 0.693 | 4.564 | 0.001 |
| t6A | OSGEPL1 | 12.223 | 0.001 | 1.331 | 0.275 |
| | TP53RK | 15.54 | 0.0006 | 1.59 | 0.179 |
| ms2i6A | CDK5RAP1 | 0.674 | 0.529 | 1.78 | 0.13 |
| m6t6A | TRMO | 0.652 | 0.54 | 1.609 | 0.174 |

**Supplemental Table 4.** PerMANOVA results for determining tRNA-modifying enzymes that exhibit altered expression in response to blood-feeding.

| Enzyme | F-statistic | P-value |
|---|---|---|
| ELP1 | 1.169 | 0.3791 |
| ELP4 | 0.5882 | 0.7094 |
| DTWD1 | 0.4563 | 0.8012 |
| OSGEPL1 | 2.001 | 0.1507 |
| TP53RK | 5.721 | 0.00631 |
| TrmO | 0.9174 | 0.5021 |
| TRMT1 | 1.64 | 0.2233 |
| TRMT44 | 1.264 | 0.3405 |
| TRMT10C | 6.244 | 0.00447 |
| TRMT5 | 6.714 | 0.00333 |
| TRMT61A1 | 8.113 | 0.0015 |
| TRMT61A2 | 4.784 | 0.01231 |
| TRMT10A | 3.766 | 0.02779 |
| TRMT13 | 7.667 | 0.00191 |
| DUS3L | 9.238 | 0.00084 |
| DUS2 | 16.27 | 5.55E-05 |
| METTL6 | 343.2 | 1.63E-12 |
| METTL2B | 11.08 | 0.00037 |
| NSUN3 | 1.681 | 0.2136 |
| TRDMT1 | 2.407 | 0.09866 |
| FTJS1A | 2.702 | 0.07353 |

| | | |
|---|---|---|
| FTJS1B | 34.71 | 9.84E-07 |
| PUS1 | 44.18 | 2.56E-07 |
| PUS3 | 1.934 | 0.162 |
| PUS7 | 21.82 | 1.21E-05 |
| PUS10 | 2.047 | 0.1436 |
| CDK5RAP1 | 1.227 | 0.3551 |
| QTRT1 | 0.3359 | 0.8816 |
| QTRT2 | 4.227 | 0.01898 |

**Supplemental Table 5**. PerMANOVA results for determining modifications affected by blood-feeding.

| Modification | F-statistic | P-value |
|---|---|---|
| Ψ | 3.263 | 0.043 |
| D | 2.774 | 0.068 |
| m3C | 2.918 | 0.059 |
| m5C | 1.238 | 0.35 |
| Cm | 1.46 | 0.272 |
| Um | 2.614 | 0.08 |
| m3U | 2.614 | 0.08 |
| I | 1.293 | 0.329 |
| Am | 1.017 | 0.449 |
| m1A | 1.742 | 0.199 |
| m2G | 1.714 | 0.205 |
| Gm | 1.047 | 0.434 |
| m1G | 13.17 | 0.0001 |
| m22G | 1.739 | 0.2003 |
| mcm5U | 5.959 | 0.005 |
| i6A | 4.069 | 0.021 |
| acp3U | 4.81 | 0.012 |
| Q | 5.206 | 0.009 |
| t6A | 4.254 | 0.018 |
| ms2i6A | 5.57 | 0.006 |
| m6t6A | 2.784 | 0.067 |
| ms2t6A | 7.319 | 0.002 |
| m7G | 2.052 | 0.142 |
| acp3D | 1.079 | 0.419 |
| m5U | 1.193 | 0.368 |
| Ψm | 4.216 | 0.019 |
| m6A | 2.735 | 0.071 |
| m1I | 2.507 | 0.089 |
| ac4C | 1.305 | 0.325 |
| m2G | 1.714 | 0.205 |
| oQ | 7.607 | 0.001 |

| | | |
|---|---|---|
| manQ | 6.599 | 0.003 |

**Supplemental Figure 3**. XIC of whole-body Tyr anticodon oligonucleotides generated by cusativin digestion. Q was not detected at position 34 as the predicted digestion product, U[Q]UAGAUCC>p (m/z = 1000) was not detected (top). An unmodified oligonucleotide was detected for the Tyr anticodon, UGUAGAUCC>p (m/z = 958) (middle). The position 34 modification on the Tyr anticodon is manQ and this was detected in the oligonucleotide U[manQ]UAGAUCC>p (m/z = 1054) (bottom). The Tyr anticodon sequence is unique and there were not any other tRNAs that contain this particular sequence, providing additional confidence in oligonucleotide identification.

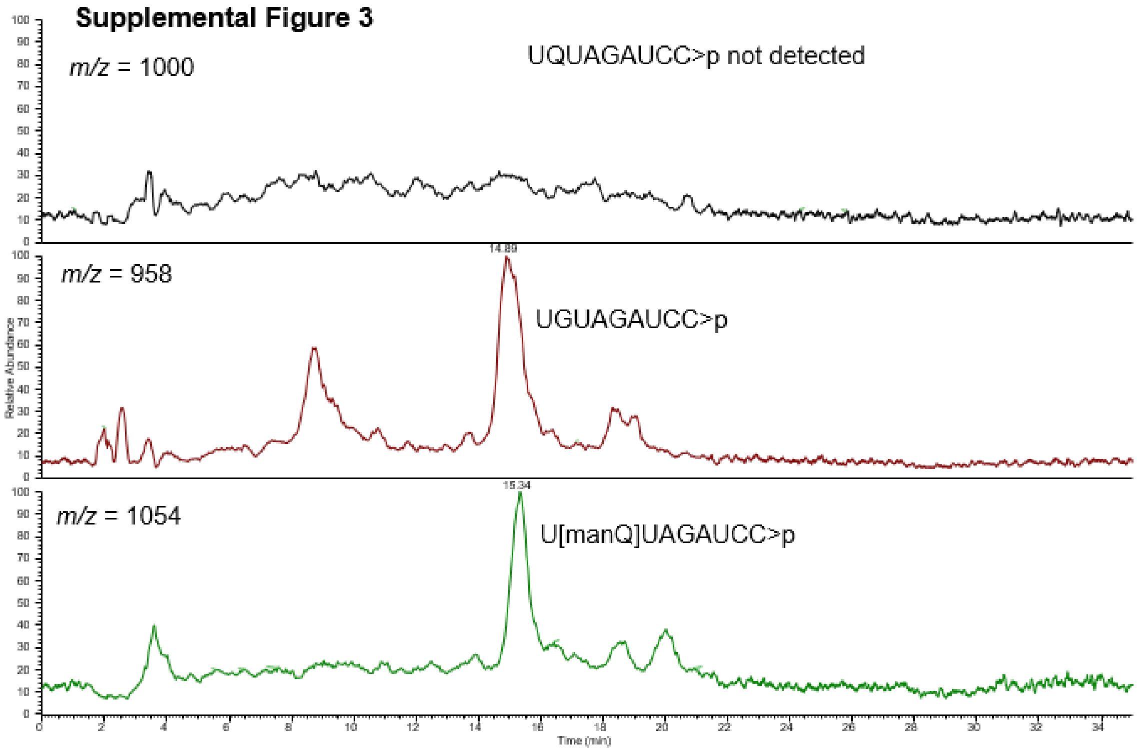
